# Effectiveness and residual activity of four common insecticides used in the Mississippi Delta to control tarnished plant bugs in cotton

**DOI:** 10.1101/2023.08.17.553780

**Authors:** Maribel Portilla, Nathan S. Little, Clint Allen, Yu Cheng Zhu

## Abstract

The tarnished plant bug, (TPB) *Lygus lineolaris* Palisot de Beauvois (Hemiptera: Miridae) is a key pest of cotton in the midsouth region and some areas of the eastern United States. Its control methods have been solely based on chemical insecticides which has contributed to insecticidal resistance and shortened residual periods for control of this insect pest. This study was conducted over a two-year period and examined the efficacy and residual effect of four commercial insecticides including lambda-cyhalothrin (pyrethroid), acephate (organophosphate), imidacloprid (neonicotinoid), and sulfoxaflor (sulfoxamine). The effectiveness and residual effects of these insecticides were determined by application on cotton field plots on four different dates during each season using three different concentrations (high: highest labeled commercial dose (CD), medium: 1/10 of the CD, low: 1/100 of the CD) on field cotton plots. Four groups of cotton leaves were randomly pulled from each treated plot and control 0-, 2-, 4-, 7-, and 9-days post treatment (DPT) and exposed to a lab colony of TPB adults. One extra leaf sample/ plot/ spray /DPT interval (0-2-4-7-9-11) during 2016 was randomly collected from the high concentration plots and sent to Mississippi State Chemical Laboratory for residual analysis. Mortality of TPB adults was greatest for those placed on leaves sprayed with the organophosphate insecticide with mortalities (%) of 81.7±23.4 and 63.3±28.8 (SE) 1-day after exposure (DAE) on leaves 0-DPT with the high concentration for 2016 and 2017, respectively, reaching 94.5±9.5 and 95.4±7.6 6-DAE each year. Mortality to all insecticides continued until 9 and 4-DPT for high and medium concentrations, respectively. However, organophosphate (39.4±28.6) and pyrethroid (24.4±9.9) exhibited higher mortality than sulfoxamine (10.6±6.6) and the neonicotinoid (4.0±1.5) 7-DAE on 9-DPT leaves with the high concentration. Based on our results using the current assay procedure, TPB adults were significantly more susceptible to contact than systemic insecticides and due to its residual effect, organophosphate could kill over 80% of the TPB population 7-DPT.

## Introduction

The tarnished plant bug, *Lygus lineolaris* Palisot de Beauvois (Hemiptera: Miridae) is the most common phytophagous species of the genus *Lygus* in North America and is widely distributed from Mexico to Alaska [1, 2]. *Lygus lineolaris* is a pest of economic importance in various agronomic crops across the United States [3]. This insect became one of the most yield limiting pests of cotton, *Gossypium hirsutum* L. (Malvales: Malvaceae) in the mid-southern U.S production system following the eradication of the boll weevil, *Anthonomus grandis* Boheman (Coleoptera: Curculionidae), and the subsequent introduction and widespread adoption of the genetically modified cotton varieties to control heliothines (collectively, the bollworm, *Helicoverpa zea* (Bodie) and tobacco budworm, *Chloridea virescens* (F.)) [3–4]. Prior to the eradication program and the introduction of transgenic cotton varieties, boll weevils and heliothines were primarily controlled with large-scale applications of organophosphates and pyrethroids (broad-spectrum insecticides). Repeated exposure to these broad-spectrum insecticides likely contributed to the development of insecticide resistance in *L. lineolaris* to several pyrethroids and some organophosphates including acephate [5–6]. The first report of *L. lineolaris* resistance to pyrethroids (bifenthrin and permethrin) was in 1993 [7], followed closely by the organophosphate methyl parathion in 1994 [8]. Both reports were from Mississippi Delta populations of *L. lineolaris*, which preceded the introduction of transgenic Bt cotton expressing insecticidal endotoxin from the soil bacterium, *Bacillus thuringiensis* (Bt) (Berliner) in the southern U.S. in 1996 [3] the start of boll weevil eradication in 1997 and predates outbreak levels of *L. lineolaris* in cotton requiring direct control [3, 7]. Since then, the number of *L. lineolaris* populations resistant to other classes of insecticides including carbamates and neoniconinoids have increased and become widespread across the southern U.S. [9–15]. The evolution of insecticide resistance in *L. lineolaris* is a major threat not only to cotton producers but to the general agricultural and public health because these increased insecticide applications due to resistance is not sustainable economically or environmentally [3, 11, 16].

Currently, multiple applications of foliar insecticides are required to adequately control *L. lineolaris* across U.S. cotton production regions [1, 17]. During the 2022 growing season, an average of 3.4 insecticide applications were being applied to control *L. lineolaris* in the U.S., which ranged from 1 application in Texas to 5.5 in the Mississippi Delta region [18]. Despite the intensity of insecticide use during this year and for the past 30 years, *Lygus* spp. continued to devastate cotton more than any other insect in all cotton producing states, except Texas (19-23). *Lygus* spp. infested a total of 2.1 million ha. in 2022 with total losses exceeding $ 324 million dollars [18]. Some studies suggested, that due to the inherent insecticidal resistance in *Lygus* spp., the residual period of insecticide activity has shortened, requiring tank mixing multiple insecticides with differing modes of action and / or two sequential applications with intervals of 4-7 d rather than two single applications [1, 17]. In general, cotton farmers around the word douse cotton crops in $ 2-3 billion worth of pesticides annually, of which $ 819 million has been classified as hazardous by the World Health Organization [24].

It has been well documented that *L. lineolaris* insecticide resistance varied between populations and among seasons [4, 11, 25, 26]. Snodgrass et al. [4] reported declines in pyrethroid resistance from the fall of one year to the spring of the following year. They mentioned that those declines are caused by the recessive nature of pyrethroid-resistance alleles and their dilution with susceptible alleles during mating. In addition, laboratory studies reported that levels of esterase, glutathione S-transferase (GST), and cytochrome P450 monoxygenase (P450) activity are directly correlated with resistance intensity to any class of insecticide used to control *L. lineolaris* [27–34]. Several authors reported elevated levels of esterase, GST, and P450 in populations of *L. lineolaris* from the Mississippi Delta and matched the highest level to the intensity of pesticide use in certain production areas [4, 102, 106]. However, a recent study demonstrated that *L. lineolaris* lost its resistance to five pyrethroids and two neonicotinoids after 36 consecutive generation under laboratory conditions without insecticide exposure. Authors mentioned that the colony lowered their activity of esterases, GST, and P450 to levels similar to the susceptible colony [35], meaning, that the insecticide resistance of *L. lineolaris* can fade away or diminishes in the absence of selection pressure. All these explained why *L. lineolaris* populations should be managed differently across seasons and various cotton production areas. They also stressed that resistance monitoring, insecticide mode of action selection, and residual effects would be vital to managing *L. lineolaris* populations as part of an effective integrated management strategy.

Today, regardless of inherent resistance levels, organophosphates and pyrethroids are the most common contact insecticides used for *L. lineolaris* control [3]. However, during the last decade newer insecticides have been introduced, including neonicotinoids and sufolxamines [2]. Therefore, this study was conducted to evaluate the efficacy and residual activity of four potential insecticides (one per class) used in Mississippi Delta. The results for this study will provide a better understanding of selecting insecticide used based in residual activity and mode of action to control *L. lineolaris* as an integrated pest management.

## Materials and methods

### Field plots and applications

The efficacy and residual activity of two contact insecticides and two systemic insecticides on *L. lineolaris* were evaluated in a non-Bt cotton cultivar in 2016 and 2017 on the Southern Insect Management Research Unit’s (SIMRU) research farm near Stoneville, MS. The experiment was set up with plots within a randomized complete block designed with four replications (4 blocks) (16 plots/block). Each plot consisted of eight rows of non-Bt cotton (DP1441RF®, Delta and Pine Land Company ^TM^, Scott, MS) with 101.6 cm wide rows approximately 100 m long. Ad hoc applications of herbicides and plant growth regulator (mepiquat chloride, Loveland Products, Inc., Morgantown, KY) were applied equally to all plots within a given year of the study. Each of the following treatments were randomly assigned to each cotton plot within each block of the experiment: 1) acephate (organophosphate) (Bracket 90 WSP^TM^, AgriSolutions, Winfield Solutions, LLC, St. Paul, MN, USA), 2) lambda-cyhalothrin (pyrethroid) (Karate^TM^, Syngenta Crop Protection, Inc. Greenboro, NC, USA), 3) imidacloprid (neonicotinoid) (Admire Pro^TM^, Bayer CropScience LP, Research Triangle Park, USA), 4) sulfoxaflor (sulfoxamine) (Transform WG^TM^, Corteva Agriscience, Indianapolis, IN, USA), and 5) untreated control. Each insecticide was applied at three different concentrations (treatments) as follow: high: highest labeled commercial dose (CD) for *L. lineolaris* in cotton, medium: 1/10 of the CD, and low: 1/100 of the CD. Each block (replication) was sprayed on four different dates within a given year of the study targeting each spray approximately two weeks apart. From the eight cotton rows/plot only the middle four rows were treated. All treatments were applied with a Lee Avenger sprayer (LeeAgra, Inc. Lubbock, TX) equipped with a ten multi-boom spray system (BellSpray Inc., Opelousa, LA) with hollow cone nozzle (TX-VS8, Tee Jet Technologies, Glendale Height, IL), which was calibrated to deliver 93.54 L of spray solution per ha. Application dates, contact and systemic insecticides used, and associated rates are listed in Table 1. Three-four-h after spray (0-day post treatment) (0-DPT), and 2, 4, 7, and 9-DPT thereafter, four groups (subsamples) of 15 leaves / group from the cotton top nodes were randomly collected from each plot (720 leaves/ sprayed block/ evaluation day) and taken to the lab for bioassays. Leaf samples were pulled from plots from the lowest to the highest concentration to avoid cross contamination.

**Table 1.** Insecticides and treatment rates for the efficacy and residual activity on *Lygus lineolaris* in cotton leaves during 2016* and 2017**.

| Insecticide | Rate | Dose/acre | Dose/hectare |
| --- | --- | --- | --- |
| Acephate | Low | 4.55 g/a | 11.2 g/ha |
|  | Medium | 45.5 g/a | 112.4 g/ha |
|  | High | 455 g/a | 1124 |
| Lambda-cyhalothrin | Low | 0.75 mL/a | 1.9 mL/ha |
|  | Medium | 7.5 mL/a | 18.5mL/ha |
|  | High | 75 mL/a | 185.3 mL/ha |
| Imidacloprid | Low | 0.5 mL/a | 1.2 mL/ha |
|  | Medium | 5 mL/a | 12.4 mL/ha |
|  | High | 50 mL/a | 123.6 mL/ha |
| Sulfoxaflor | Low | 0.65 g/a | 1.6 g/ha |
|  | Medium | 6.5 g/a | 16.1 g/ha |
|  | High | 65 g/a | 160.6 g/ha |

Application dates:
\*2016: July 12, July 20, July 25, and August 8
\*\*2017: July 17, July 30, August 21, and Sep 6

### Leaf samples for insecticide residual analysis test

From at least two sprays, each insecticide had leaf samples pulled at 0-day post treatment (0-DPT), 2, 4, 7, 9, and 11-DPT. The top of 10 cotton plants (high concentration plots) were marked in each plot with flagging tape just before the plots were sprayed. One leaf from each flagged plant was pulled (10 leaves/sample/plot/insecticide) at each evaluation time. Leaf samples were sent to the Mississippi State University Chemistry Laboratory for leaves that were present at the time of the insecticide spray. The insecticide analysis test was done for 2016 only.

### Insect colonies

A laboratory-reared *L. lineolaris* colony has been maintained at the United States Department of Agriculture Agricultural Research Service (USDA ARS), Southern Insect Management Research Unit (SIMRU) in Stoneville, MS since 2011 [36], but was stablished in 1998 at the USDA-ARS Biological Control Rearing and Research Unit in Starkville, MS [37]. This laboratory colony is routinely reared following procedures outlined by Portilla et al. [37], which was designed for mass production of even-aged individuals. Insects were held in environmental chambers with a photoperiod of 12:12 (L: D) h, 27C, and 60% RH. Mixed-sex adults 2-d old were used for this study. This insect colony has not been exposed to insecticides and/or have had field population infusions, which make the insects particularly valuable for screening insecticides.

### Bioassay procedure

Laboratory essays were conducted to determine the efficacy and residual activity of the insecticide treatments. Four groups of 15 leaves randomly collected from each plot at 0 (the day of the spray) (60 leaves / plot) (960 leaves / block / evaluation time) were taken to the lab and placed individually into 30-mL plastic cups (T-125 SOLO-cup, Pleasant Prairie, WI). A 2-d old *L. lineolaris* adult (unknown sex) was released to each cup that contained the treated leaf with the three concentrations of each tested insecticide. The lids for the cups had three holes for ventilation (3 mm diameter). Adults were examined daily (7-d period) for mortality. This process was repeated for each evaluation time at 0, 2, 4, 7, and 9-DPT and for each spray (replication) (4,800 leaves-insects/spray-replicate).

### Analyses

All experiments were analyzed using SAS 9.4 (SAS Institute 213) [38]. A randomized complete block design with factorial arrangements (treatment concentration-days of exposure x subsamples x spray) (32 x 4 x 4) per each evaluation time was used for cumulative mortality. Mortality percentage and residual analysis test were analyzed by using ANOVA followed by Tukey’s HSD. Probit analyses were used to develop regressions for estimating lethal concentrations (LC_50_) of each tested insecticide (SAS Institute 2013). Percent mortalities for each treatment were corrected for control effects using Abbott’s formula [39]. Resistance ratios (RR_50_) and 95% CI were calculated using Robertson and Priestler’s formula [40]. Bioassays were considered significant when the slope of the line was significant (*P* < 0.05).

## Results

### Effectivity and residual activity of Acephate by leaf sample bioassay

There were statistically significant differences in cumulative mortality of acephate (organophosphate) to *L. lineolaris* among concentrations and day of exposure at each evaluation of DPT (Table 2). The organophosphate had the greatest effect on TPB adults with mortality (%) of 48.75 ± 18.3 0-day after exposure (DAE) on leaves 0-DPT and its residual effect continued in leaves pulled 9-DAT. The high residual activity was sustained until 4-DPT period with mortality over 90% on 7-DAE for 2016 (Figs 1ABC), decreasing then to 80 ± 28.8 on 7-DAT (Fig 1D) and 39 ± 8.27 on 9-DAT (Fig 1E). A similar cumulative mortality trend was observed only for the high concentration in 2017 with no significant differences on cumulative mortality from 4 to 7-DAE for both years (Figs 1AF). The high concentration had a better control in 2016, while a better dose response was observed in 2017 with a greater residual activity for the lower concentrations (Figs 1FGHI). Heavy precipitation affected residual activity in 2017 for the 7-DPT (Fig 1I), lowering its residual effect 5.14-fold (12.08 ± 2.93) compared to the cumulative mortality at 4-DPT (62.08 ± 34.74) or 15-fold lower if compared with the 7-DPT evaluation in 2016. Due to its low residual activity, no leaf samples were collected for the 9-DPT evaluation in 2017. Less than 2% cumulative mortality was observed in control with no significant differences among medium and low concentrations in 2016 for 4, 7, and 9-DPT (Figs 1CDE) and in 2017 for 7-DPT only (Fig 1I). Table 3 shows that the probit model produced a good fit of the data for the LC_50_ estimated values for both years. The residual activity is measured by the estimation of the LC_50_s and resistance ratios (RR50s). The lethal concentration and RR_50_s for 2016 and 2017 increased significantly over the time (DPT). Therefore, the lower the residual activity the higher LC_50_s and RR_50_s.

**Table 2.** Overall General Lineal Model for cumulative mortality (7-d period) of insecticides and treatments rates on *Lygus lineolaris* in cotton leaves at different days post treatment during 2016 and 2017.

| Insecticide | Days Post Treatment (DPT) |  |  |  |  |
| --- | --- | --- | --- | --- | --- |
|  | 0-D | 2-D | 4-D | 7-D | 9-D |
| 2016 |  |  |  |  |  |
| Acephate | F = 214.2<br>*P* = <.0001 | F = 239.45<br>*P* = <.0001 | F = 159.25<br>*P* = <.0001 | F = 8.67<br>*P* = <.0001 | F = 14.39<br>*P* = <.0001 |
| Lambda-cyhalothrin | F = 21.93<br>*P* = <.0001 | F = 90.19<br>*P* = <.0001 | F = 67.60<br>*P* = <.0001 | F = 16.57<br>*P* = <.0001 | F = 21.16<br>*P* = <.0001 |
| Imidacloprid | F = 23.79<br>*P* = <.0001 | F = 7.70<br>*P* = <.0001 | F = 9.95<br>*P* = <.0001 | F = 13.18<br>*P* = <.0001 | F = 4.67<br>*P* = <.0001 |
| Sulfoxaflor | F = 78.69<br>*P* = <.0001 | F = 38.09<br>*P* = <.0001 | F = 32.01<br>*P* = <.0001 | F = 8.67<br>*P* = <.0001 | F = 10.37<br>*P* = <.0001 |
| 2017 |  |  |  |  |  |
| Acephate | F = 107.09<br>*P* = <.0001 | F = 79.76<br>*P* = <.0001 | F = 26.20<br>*P* = <.0001 | F = 10.94<br>*P* = <.0001 |  |
| Lambda-cyhalothrin | F = 45.97<br>*P* = <.0001 | F = 45.24<br>*P* = <.0001 | F = 24.33<br>*P* = <.0001 | F = 11.99<br>*P* = <.0001 |  |
| Imidacloprid | F = 14.06<br>*P* = <.0001 | F = 8.69<br>*P* = <.0001 | F = 7.07<br>*P* = <.0001 | F = 5.04<br>*P* = <.0001 |  |
| Sulfoxaflor | F = 79.76<br>*P* = <.0001 | F = 7.89<br>*P* = <.0001 | F = 7.49<br>*P* = <.0001 | F = 6.87<br>*P* = <.0001 |  |

n = 60
DF = 31, 3 for all treatments at 0, 2, and 4 DPT 2016
DF = 31, 2 for all treatments at 7 and 9 DPT 2017
DF = 31, 3 for all treatments at 0, 2, 4, and 7 DPT 2017

**Figure 1.**
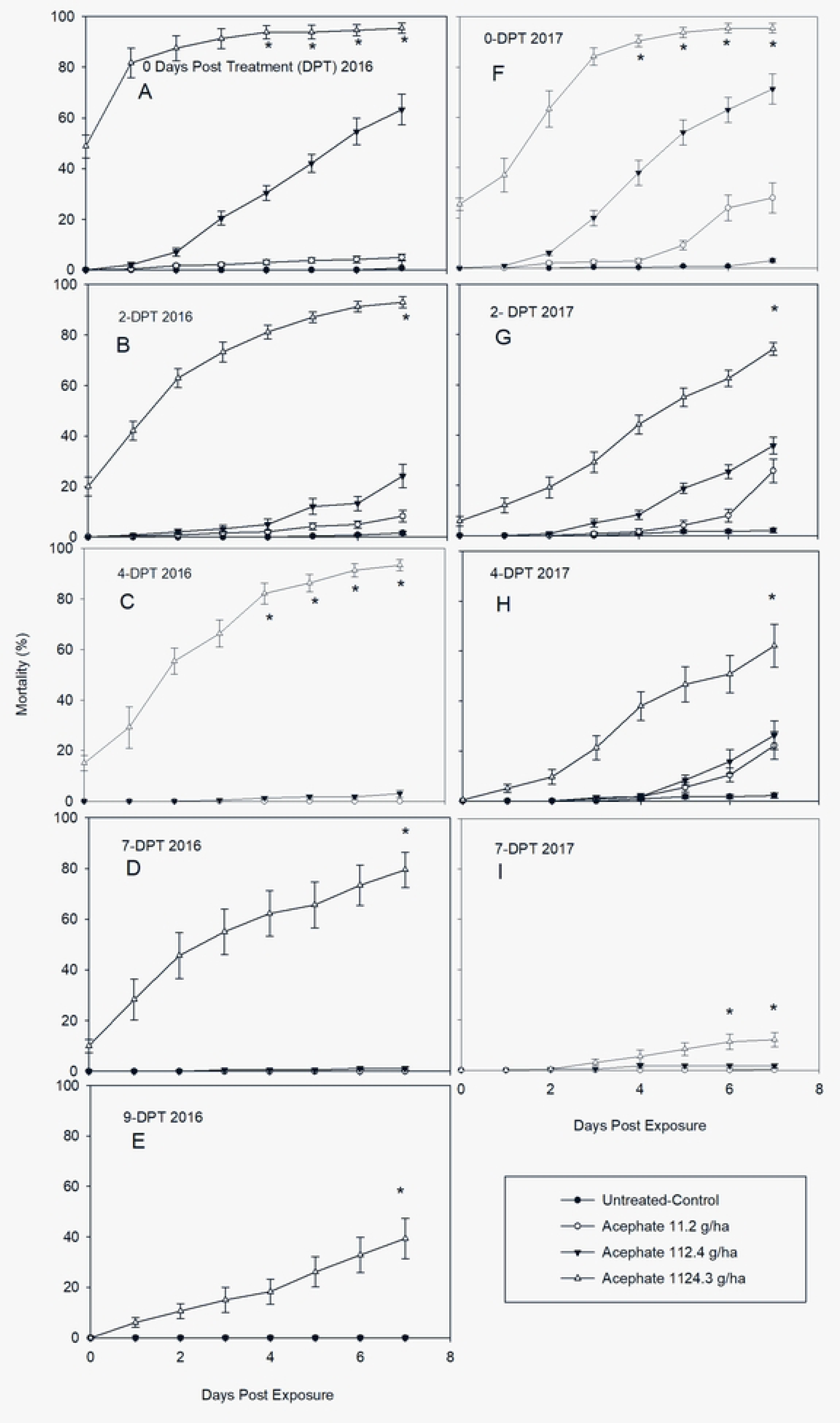
Post treatment mean ± SD cumulative mortality of a laboratory-reared *Lygus lineolaris* colony exposed to acephate on cotton leaves sprayed in the field and pulled days posttreatment (DPT). A-E: Mortality (%) in 2016 after DPT period. F-I: Mortality (%) in 2017 after DPT periods (Tukey’ HSD test, *p* = 0.05).

**Table 3.** Lethal mortality response (LC_50_) of *Lygus lineolaris* exposed to different concentrations of acephate estimated at different days post treatment on cotton leaves during 2016 and 2017.

| Days Post Treatment | Concentration response (g / ha) |  |  |  |  |  |  |  |
| --- | --- | --- | --- | --- | --- | --- | --- | --- |
|  | n | Slope ± SE | LC<sub>50</sub> (95%CI) | Probit Trend |  |  |  | RR<sub>50</sub> (95%CI) |
|  |  |  |  | Test for Slope |  | Test for GoF |  |  |
|  |  |  |  | X<sup>2</sup> | P > X<sup>2</sup> | X<sup>2</sup> | P > X<sup>2</sup> |  |
| Year 2016 |  |  |  |  |  |  |  |  |
| 0-D | 960 | 0.743 ± 0.076 | 86.12 (63.18 - 115.1) | 96.320 | <.0001 | 2.8432 | <.0001 | 1.0 |
| 2-D | 960 | 0.980 ± 0.122 | 256.71 (192 - 342.20) | 64.592 | <.0001 | 2.6776 | <.0001 | 2.86 (1.9 – 4.29) |
| 4-D | 960 | 1.292 ± 0.108 | 283.18 (230.1 - 346.44) | 142.501 | <.0001 | 1.8312 | 0.0005 | 3.36 (2.33 – 4.78) |
| 7-D | 720 | 1.133 ± 0.153 | 414.15 (289.06 - 568.1) | 54.310 | <.0001 | 3.4325 | <.0001 | 4.84 (3.13 – 7.49) |
| 9-D | 720 | 0.591 ± 0.102 | 839.86 (525.6 - 1627.2) | 33.781 | <.0001 | 3.0014 | <.0001 | 9.87 (5.45 – 17-84) |
| Year 2017 |  |  |  |  |  |  |  |  |
| 0-D | 960 | 0.511 ± 0.067 | 40.06 (23.7 - 62.34) | 58.461 | <.0001 | 3.4623 | <.0001 | 1.0 |
| 2-D | 960 | 0.290 ± 0.039 | 200.65 (120.8 - 349.65) | 56.267 | <.0001 | 1.8675 | 0.0003 | 4.89 (2.46 – 9.74) |
| 4-D | 960 | 0.252 ± 0.689 | 582.92 (205.5 - 4502.3) | 13.531 | 0.0002 | 5.5291 | <.0001 | 13.99 (3.92 – 49.96) |
| 7-D | 960 | 0.464 ± 0.1001 | 14615 (5621-145339) | 20.903 | <.0001 | 1.1410 | 0.2349 | 391.65 (92.54 – 1657.44) |

LC<sub>50</sub> values were calculated in g / ha Mortality was scored at 7 Days After Exposure Differences among RR<sub>50</sub> values are significant if 95% CI do not include 1.0 RR<sub>50</sub> and 95% CI calculated using formula from Robertson et al. [40]

### Effectivity and residual activity of lambda-cyhalothrin by leaf sample bioassay

As observed with acephate, there were statistically significant differences in cumulative mortality of lambda-cyhalothrin (pyrethroid) to *L. lineolaris* among concentrations and day of exposure at each evaluation period of DPT (Table 2). However, the pyrethroid had lower effect relative to the organophosphate on TPB adults. Although its residual effect continued to 9-DPT for the high and medium concentration, its initial mortality on 0-DAE did not exceed 10% for 2016 (8 ± 1.6) but increased over 20% in 2017 (22.50 ± 6.85) on leaves pulled on 0-DPT for the high concentration (Figs 2A and F). Fig 2A and F show that initial mortality for the medium concentration started 1-DAE, while for the low concentration, started 5-DAE for both years on leaves pulled on 0-DPT. Cumulative mortality for the high concentration slowly reached about 80% 7-DAE (Figs 2A and F). This residual activity continued in leaves pulled 2-DPT (79.17 ± 13.30) (Fig 2B) and 4-DPT (69.58 ± 22.43) (Fig 2C) for 2016, and 2-DPT (75.0 ± 23.28) for 2017 (Fig 2G). Cumulative mortality for the medium concentration reached 35.41 ± 8.62 and 55.0 ± 7.56 for 2016 and 2017, respectively 7-DAE on leaves 0-DPT (Figs 2A and F). Its residual activity dropped 2.9-(12.50 ± 2.34) and 2.8-fold (20.83 ± 6.71) in 2016 and 2017, respectively on 2-DPT, lowering to less than 5.0 and 2.0 % on 4-DPT for 2016 and 2017, respectively (Figs 2B and G). No residual activity was observed for the low concentration after 2-DPT for either year (Figs 2C and H). Similar mortality trends were observed for both years during the evaluation periods of 0, 2, and 4-DPT. However, similar to that observed for acephate, the residual activity of lambda-cyhalothrin was likely lessened due to the high precipitation event before the 7-DPT on 2017 (Fig 2I). Due to its low residual activity, no leaf samples were collected for the 9-DPT evaluation in 2017. Less than 2% cumulative mortality was observed in the control with no significant differences among low concentrations in 2016 for 4, 7, and 9-DPT (Fig 2CDE) and in 2017 for 4, and 7-DPT (Figs 1H and I). Table 3 shows a dose-dependent effect at all evaluation period (DPT) for both years. The LC_50_ value for leaves pulled 0-DPT was 33.85 mL / ha, lowering its residual activity 10.59-fold after 9 days (358.79.2 mL / ha) for 2016. In 2017 the LC_50_ for 0-DPT was 15.1 mL / ha decreasing the residual activity 17.65-folds (254.1 mL / ha) after 7 days. Probit model produced a good fit of the data for the LC_50_ estimated values for both years.

**Figure 2.**
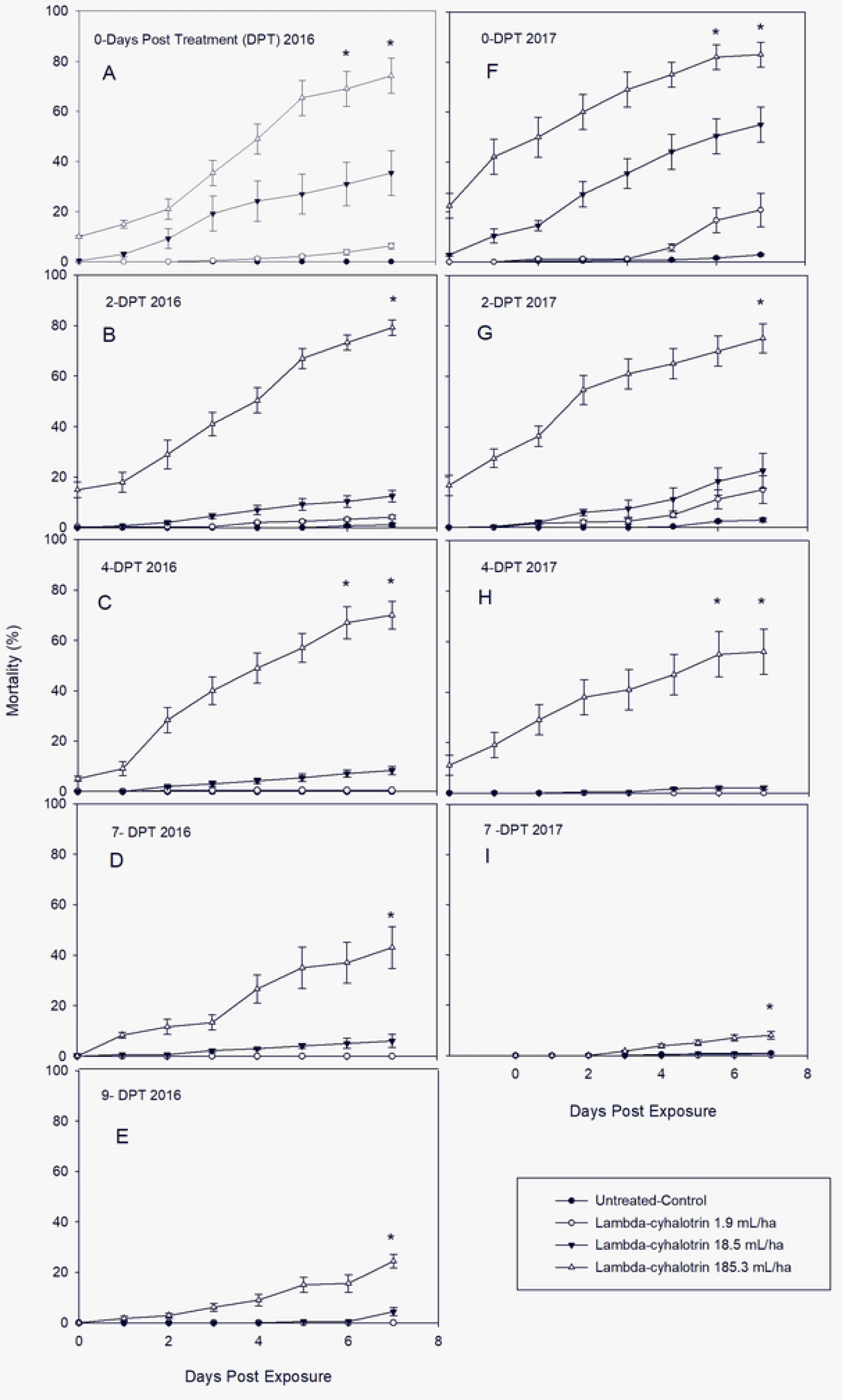
Post treatment mean ± SD cumulative mortality of laboratory-reared *Lygus lineolaris* colony exposed to the pyrethroid insecticide lambda-cyhalothrin on cotton leaves sprayed in the field and pulled days posttreatment (DPT). A-E: Mortality (%) in 2016 after DPT periods. F-I: Mortality (%) in 2017 after DPT periods (Tukey’ HSD test, *p* = 0.05).

### Effectivity and residual activity of imidacloprid by leaf sample bioassay

The effectiveness and residual activity of the neonicotinoid imidacloprid, is displayed in Fig 3. There were statistically significant differences in cumulative mortality of imidacloprid to *L. lineolaris* among concentrations and day of exposure at each evaluation period of DPT (Table 2). The efficacy of the neonicotinoid against *L. lineolaris* was distinctly inferior over organophosphate and pyrethroid in terms of mortality percentage. Although the residual effect continued to 9-DPT for the high concentration, the cumulative mortality (7-DAE) barely exceeded 30% for both years (33.33 ± 12.17 and 29.58 ± 19,77 for 2016 and 2017, respectively) on leaves pulled 0-DPT. The cumulative mortality decreased > 4-fold for 2016 (8.33 ± 3.07) but it was sustained at > 20% for 2017 (22.08 ± 7.7) on leaves pulled 2-DPT (Figs 3B and G). After that, the residual effect was not enough to kill more than 20% of the population in 2016 until 7-DPT period (Figs 3BC and D) and until 4-DPT period for 2017 (Figs 3G and H). The initial mortality of TPB adults with the highest concentration was observed 1 and 2-DAE on leaves pulled 0-DPT for 2016 and 2017, respectively. The initial mortality for the same treatment was exhibited 2 and 5-DAE for 2016 and 2017, respectively on leaves pulled 2-DPT period (Figs 3B and G). No differences in mortality were observed between control, medium, and low concentrations at 4, 7, and 9-DPT for 2016 (Figs 3CD and E) and 4 and 7-DPT for 2017 (Figs 3H and I). Mortality in untreated leaves did not occur until 6-7 DAE. Although the efficacy and residual activity was ineffective for the beginning of the treatment applications, it was very clear how the high precipitation obtained in 2017 also affected this insecticide (Fig 3I). Table 5 shows a dose-dependent effect at all evaluation periods (DPT) for 2016. The probit model did not produce a good fit for all data regarding LC_50_ estimated values for 2017. The LC_50_ value for leaves pulled 0-DPT in 2016 was 421.6 mL / ha, that was 3.4-fold higher that the commercial dose (123.6 mL / ha) decreasing the residual activity 11.71-fold (5505.5 mL / ha) after 9-DPT. An unexpected irregularity of LC_50_ estimation was observed for 2017; therefore, the residual activity could not be measured by the estimation of LC_50_s and RR_50_s for this year. However, Fig 5 clearly demonstrated that the efficacy gradually decreased throughout the DPT periods.

**Figure 3.**
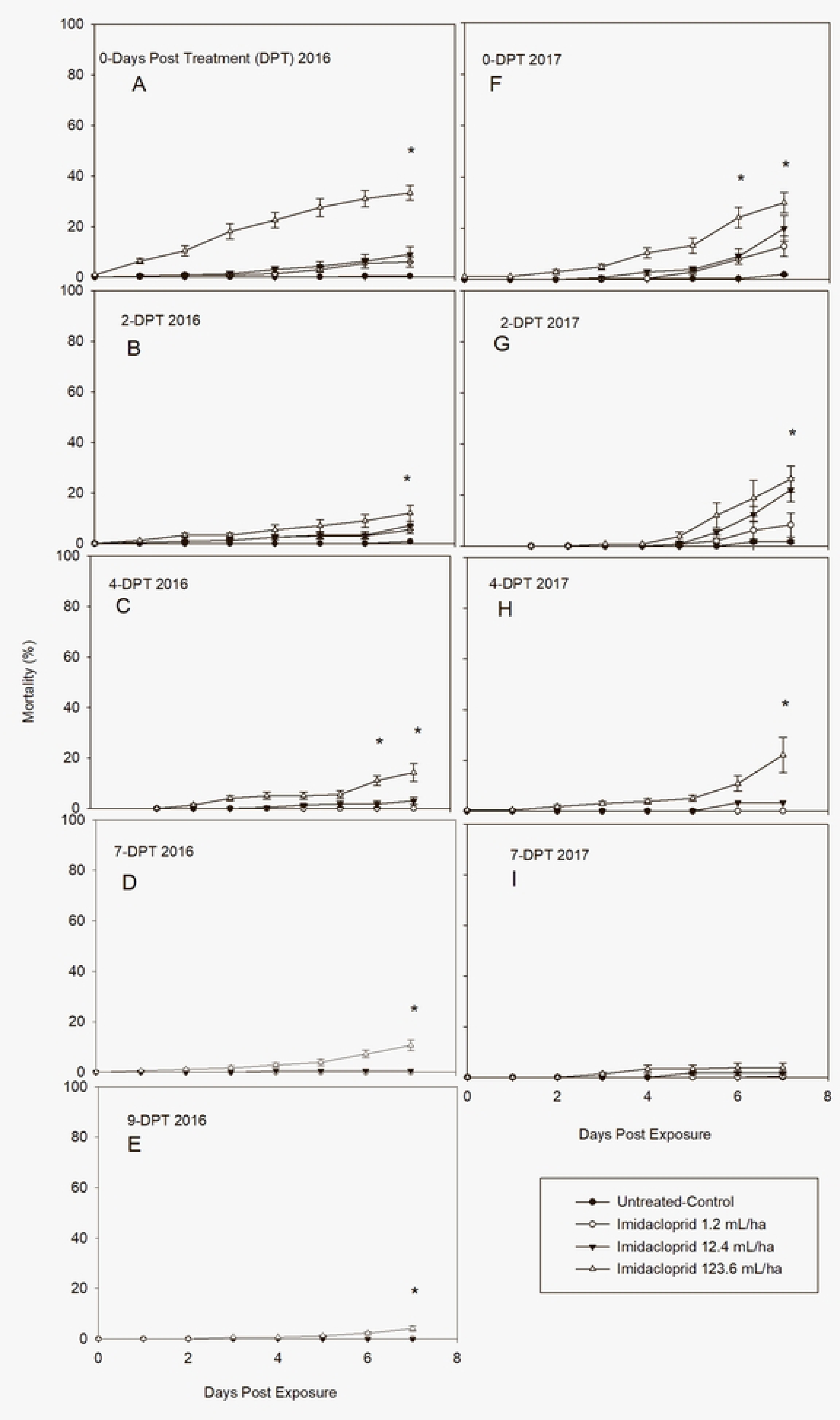
Post treatment mean ± SD cumulative mortality of a laboratory-reared *Lygus lineolaris* colony exposed to the insecticide neonicotinoid, imidacloprid, on cotton leaves sprayed in the field and pulled days posttreatment (DPT). A-E: Mortality (%) in 2016 after DPT periods. F-I: Mortality (%) in 2017 after DPT periods (Tukey’ HSD test, *p* = 0.05).

**Table 4.** Lethal mortality response (LC_50_) of *Lygus lineolaris* exposed to different concentrations of lambda-cyhalothrin estimated at different days post treatment in cotton leaves during 2016 and 2017.

| Days Post Treatment | Concentration response (mL / ha) |  |  |  |  |  |  |  |
| --- | --- | --- | --- | --- | --- | --- | --- | --- |
|  | n | Slope ± SE | LC<sub>50</sub> (95%CI) | Probit Trend |  |  |  | RR<sub>50</sub> (95%CI) |
|  |  |  |  | Test for Slope |  | Test for GoF |  |  |
|  |  |  |  | X<sup>2</sup> | P > X<sup>2</sup> | X<sup>2</sup> | P > X<sup>2</sup> |  |
| Year 2016 |  |  |  |  |  |  |  |  |
| 0-D | 960 | 0.340 ± 0.054 | 33.85 (14.21 - 77.34) | 38.895 | <.0001 | 3.3980 | <.0001 | 1 |
| 2-D | 960 | 0.895 ± 0.084 | 61.65 (42.28 - 94.15) | 112.569 | <.0001 | 1.4243 | 0.0307 | 1.81 (0.75 – 4.41) |
| 4-D | 960 | 0.539 ± 0.058 | 76.58 (61.75 - 93.9) | 85.057 | <.0001 | 2.3410 | <.0001 | 2.17 (0.95 – 4.95) |
| 7-D | 720 | 0.501 ± 0.081 | 126.52 (72 -271.57) | 37.802 | <.0001 | 2.9890 | <.0001 | 3.73 (1.36 – 10.18) |
| 9-D | 720 | 0.406 ± 0.049 | 358.8 (212.76 - 742.2) | 68.614 | <.0001 | 1.2139 | 0.1826 | 10.59 (3.89 – 28.81) |
| Year 2017 |  |  |  |  |  |  |  |  |
| 0-D | 960 | 2.984 ± 9711 | 15.07 (-) | 0.0000 | 0.9998 | 1.2604 | 0.2467 | 1 |
| 2-D | 960 | 0.467 ± 0.552 | 36.42 (-) | 0.715 | 0.3979 | 1.7594 | 0.0136 | 1.3 (-) |
| 4-D | 960 | 0.550 ± 1.115 | 71.96 (35.68 – 201.1) | 24.301 | <.0001 | 0.7068 | 0.9051 | 5.22 (-) |
| 7-D | 960 | 0.750 ± 0.289 | 254.83 (147.9 - 1662) | 6.708 | 0.0096 | 0.8077 | 0.8177 | 17.65 (-) |

LC<sub>50</sub> values were calculated in mL / ha
Mortality was scored at 7 Days After Exposure
Differences among RR<sub>50</sub> values are significant if 95% CI do not Include 1.0
RR50 and 95% CI calculated using formula from Robertson et al. [40]

**Table 5.** Lethal mortality response (LC_50_) of *Lygus lineolaris* exposed to different concentrations of imidacloprid estimated at different days post treatment in cotton leaves during 2016 and 2017.

| Days Post Treatment | Concentration response (g – mL/Ha) |  |  |  |  |  |  |  |
| --- | --- | --- | --- | --- | --- | --- | --- | --- |
|  | n | Slope ± SE | LC<sub>50</sub> (95%CI) | Probit Trend |  |  |  | RR<sub>50</sub> (95%CI) |
|  |  |  |  | Test for Slope |  | Test for GoF |  |  |
|  |  |  |  | X<sup>2</sup> | P > X<sup>2</sup> | X<sup>2</sup> | P > X<sup>2</sup> |  |
| Year 2016 |  |  |  |  |  |  |  |  |
| 0-D | 960 | 0.263 ± 0.066 | 421.6 (216.5 – 1215.2) | 15.910 | <.0001 | 1.8262 | 0.0005 | 1 |
| 2-D | 960 | 0.604 ± 0.268 | 456.7 (224.6 - 41175016) | 5.083 | 0.0242 | 1.5543 | 0.0094 | 1.09 (0.20 – 5.89) |
| 4-D | 960 | 0.368 ± 0.051 | 470.5 (184.6 - 3301) | 53.332 | <.0001 | 1.3355 | 0.0636 | 1.11 (0.27 – 4.56) |
| 7-D | 720 | 0.328 ± 0.061 | 798.9 (289.8 - 6311) | 28.740 | <.0001 | 1.4018 | 0.0601 | 1.69 (0.29 – 9.82) |
| 9-D | 720 | 0.255 ± 0.055 | 5505.5 (1058.6 - 283783) | 21.086 | <.0001 | 1.1363 | 0.2682 | 11.71 (0.85 – 161.18) |
| Year 2017 |  |  |  |  |  |  |  |  |
| 0-D | 960 | 0.135 ± 0.063 | 9350 (365.9 - 26788) | 4.702 | 0.0301 | 3.8583 | <.0001 | 34.92 (0.10 -11726.55) |
| 2-D | 960 | 0.029 ± 0.084 | 6.63472E12 (-) | 0.119 | 0.7294 | 8.2031 | <.0001 | (-) |
| 4-D | 960 | 0.425 ± 0.082 | 219.9 (105.3 - 832.5) | 26.770 | <.0001 | 3.5297 | <.0001 | 1 |
| 7-D | 960 | 0.252 ± 0.114 | 145441 (2713 - 6.74672E37) | 4.921 | 0.0265 | 1.3444 | 0.0593 | 18.69 (0.51 – 843292) |

LC<sub>50</sub> values were calculated in g/Ha Mortality was scored at 7 Days After Exposure Differences among RR<sub>50</sub> values are significant if 95% CI do not include 1.0 RR50 and 95% CI calculated using formula from Robertson et al. [40]

### Effectivity and residual activity of sulfoxaflor by leaf sample bioassay

The effectiveness and residual activity of the sulfoxaflor (sulfoxamine) is presented in Fig 3. There were statistically significant differences in cumulative mortality of sulfolxaflor to *L. lineolaris* among concentrations and day of exposure at each evaluation period of DPT (Table 2). Like all insecticides tested, the residual effect of the high concentration continued until after 9-DPT for 2016 (Fig 4E) and 7-DPT for 2017 (Fig 4I). The mortality trend for sulfoxaflor is comparable to pyrethroid for the leaves pulled 0-DPT, where mortality increased gradually and reached the maximum cumulative mortality 7-DAE, for the high concentration for both years (67.91 ± 15.62 and 74.16 ± 10.0 for 2016, 2017, respectively) (Figs 4A and F). However, after the 2-DPT the residual effect decreased almost as fast as the neonicotinoid either in 2016 or 2017 (Fig 4GHI). The initial mortality for the high concentration was observed 2-DAE in 2016, while in 2017 a low percentage (5.83 ± 2.42) was observed at 0-DPT. No differences in mortality were observed between control, medium, and low concentrations at 2, 4, 7, and 9-DPT for 2016 (Figs 4BCD and E) and 4 and 7-DPT for 2017 (Figs 3H and I). Mortality in untreated leaves did not occur until 6-7 DAE. Similarly, as the rest of the insecticides, sulfoxaflor was affected by the precipitation in 2017. Table 6 shows a dose-dependent effect at all evaluation period (DPT) for 2016 and 2017. Probit model did produce a good fit for all data for the LC_50_ estimated values for both years. The LC_50_ value for leaves pulled 0-DPT was 91.23 g / ha, lowering its residual activity 20.48-fold after 9 days (1763.3 g / ha) for 2016. In 2017 the LC_50_ for 0-DPT was 156.96 g / ha decreasing the residual activity 434.53-folds (254.1 g / ha) after 7 days. Probit model produced a good fit of the data for the LC_50_ estimated values for both years except for the 2-DPT in 2017 which was higher than expected.

**Figure 4.**
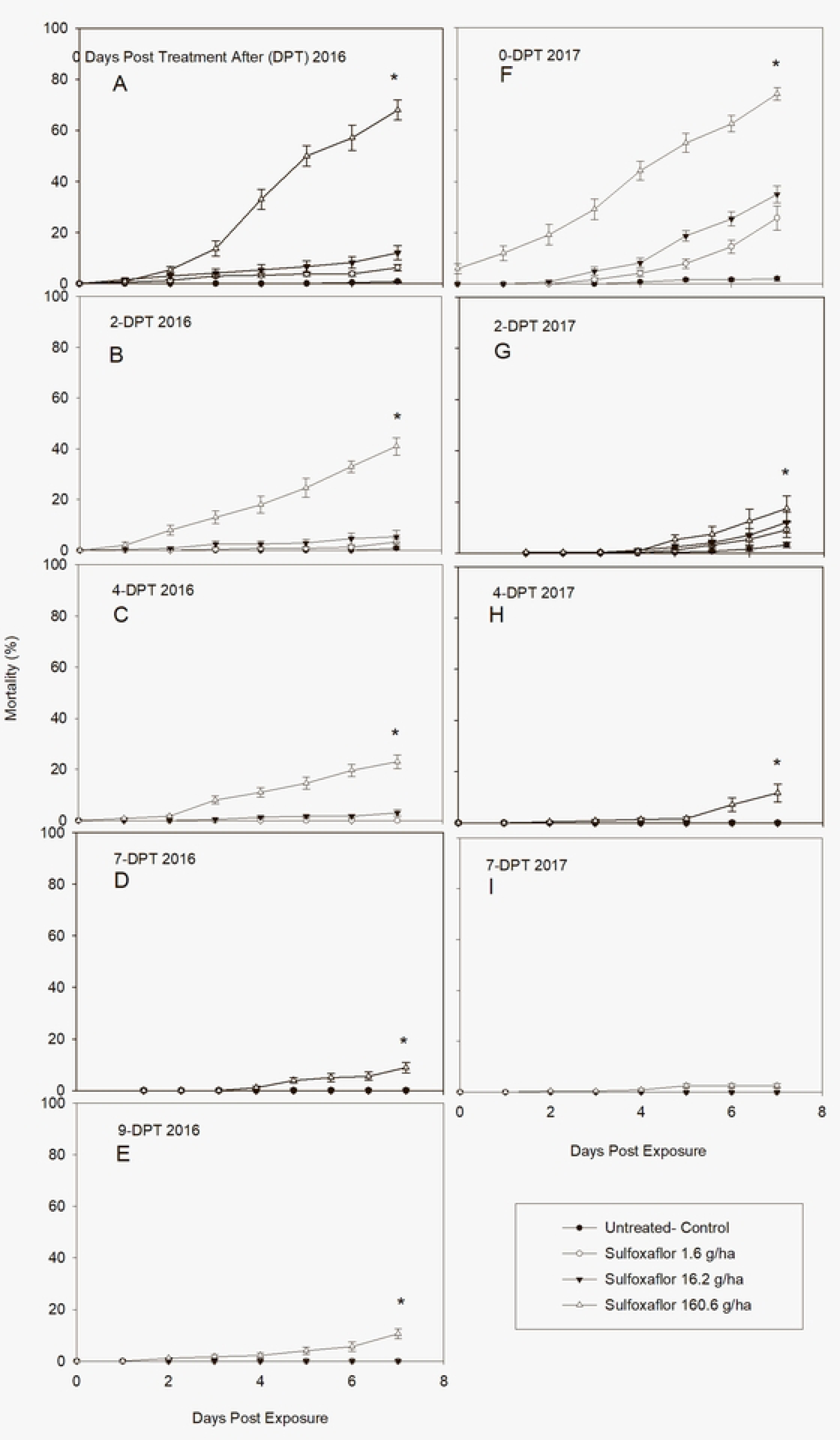
Post treatment mean ± SD cumulative mortality of a laboratory-reared *Lygus lineolaris* colony exposed to the sulfoxamine insecticide sulfoxaflor, on cotton leaves sprayed in the field and pulled days posttreatment (DPT). A-E: Mortality (%) in 2016 after DPT periods. F-I: Mortality (%) in 2017 after DPT periods (Tukey’ HSD test, *p* = 0.05).

**Table 6.** Lethal mortality response (LC_50_) of *Lygus lineolaris* exposed to different concentrations of sulfoxaflor estimated at different days post treatment in cotton leaves during 2016 and 2017.

| Days Post Treatment | Concentration response (g / ha) |  |  |  |  |  |  |  |
| --- | --- | --- | --- | --- | --- | --- | --- | --- |
|  | n | Slope ± SE | LC<sub>50</sub> (95%CI) | Probit Trend |  |  |  | RR<sub>50</sub> (95%CI) |
|  |  |  |  | Test for Slope |  | Test for GoF |  |  |
|  |  |  |  | X<sup>2</sup> | P > X<sup>2</sup> | X<sup>2</sup> | P > X<sup>2</sup> |  |
| Year 2016 |  |  |  |  |  |  |  |  |
| 0-D | 960 | 0.758 ± 0.104 | 91.23 (69.39 – 119.6) | 53.710 | <.0001 | 1.8460 | 0.0004 | 1 |
| 2-D | 960 | 0.667 ± 0.158 | 328.9 (162.10 – 520.2) | 17.871 | <.0001 | 2.2846 | <.0001 | 2.67 (1.56 – 4.56) |
| 4-D | 960 | 0.445 ± 0.056 | 261.7 (161.60 – 520.9) | 64.162 | <.0001 | 1.5275 | 0.0121 | 3.04 (1.65 – 5.61) |
| 7-D | 720 | 0.287 ± 0.061 | 844.8 (339.26 – 5201.6) | 22.183 | <.0001 | 1.4790 | 0.0356 | 9.81 (2.94 – 32.77) |
| 9-D | 720 | 0.357 ± 0.065 | 1763.3 (483.60 - 36507) | 30.04 | <.0001 | 1.4000 | 0.0608 | 20.48 (3.46 – 121.12) |
| Year 2017 |  |  |  |  |  |  |  |  |
| 0-D | 960 | 0.254 ± 0.070 | 156.9 (52.31- 2183) | 13.034 | 0.0003 | 5.3977 | <.0001 | 1 |
| 2-D | 960 | 0.110 ± 0.082 | 2284306 (-) | 1.801 | 0.1796 | 3.6038 | <.0001 | 10744 (0.001 – 1043569415) |
| 4-D | 960 | 0.290 ± 0.057 | 1287 (406.8 - 15201) | 25.803 | <.0001 | 1.8681 | 0.0003 | 8.95 (1.14 – 70.29) |
| 7-D | 960 | 0.199 ± 0.049 | 62473 (4961 - 77476623) | 16.342 | <.0001 | 0.8597 | 0.7378 | 434.53 (7.72 – 24472.75) |

LC<sub>50</sub> values were calculated in g / ha Mortality was scored at 7 Days After Exposure Differences among RR<sub>50</sub> values are significant if 95% CI do not include 1.0 RR50 and 95% CI calculated using formula from Robertson et al. [40]

### Leaf samples for insecticide residual analysis test

Timing of leaf samples and the residual analysis test is presented in Fig 5. Residual activity of all insecticides was detected up to 11-DPT. There were statistically significant differences in residual activity (ppm) among insecticides and period of evaluations 0, 2, 4, and 7-DPT. Acephate exhibited the greatest residual effect with the highest ppm values. It was highly significantly different among lambda cyhalothrin, imidacloprid, and sulfoxaflor on leaves collected 0-DPT: F _(3, 3)_ = 10.22, *P* = 0.0060; 2-DPT: F _(3, 3)_ = 12.31, *P* = 0.0057; 4-DPT: F _(3, 3)_ = 8.15, *P* = 0.0593; 7-DPT: F _(3, 3)_ = 21.33, *P* = 0.0159; and 9-DPT: F _(3, 3)_ = 21.33, *P* = 0.0357. No significant differences were observed between lambda cyhalothrin, imidacloprid, and sulfoxaflor at any of those evaluation times. No significant differences of residual activity (ppm) were found among insecticides for 11-DPT (*P* = 0.5692).

**Figure 5.**
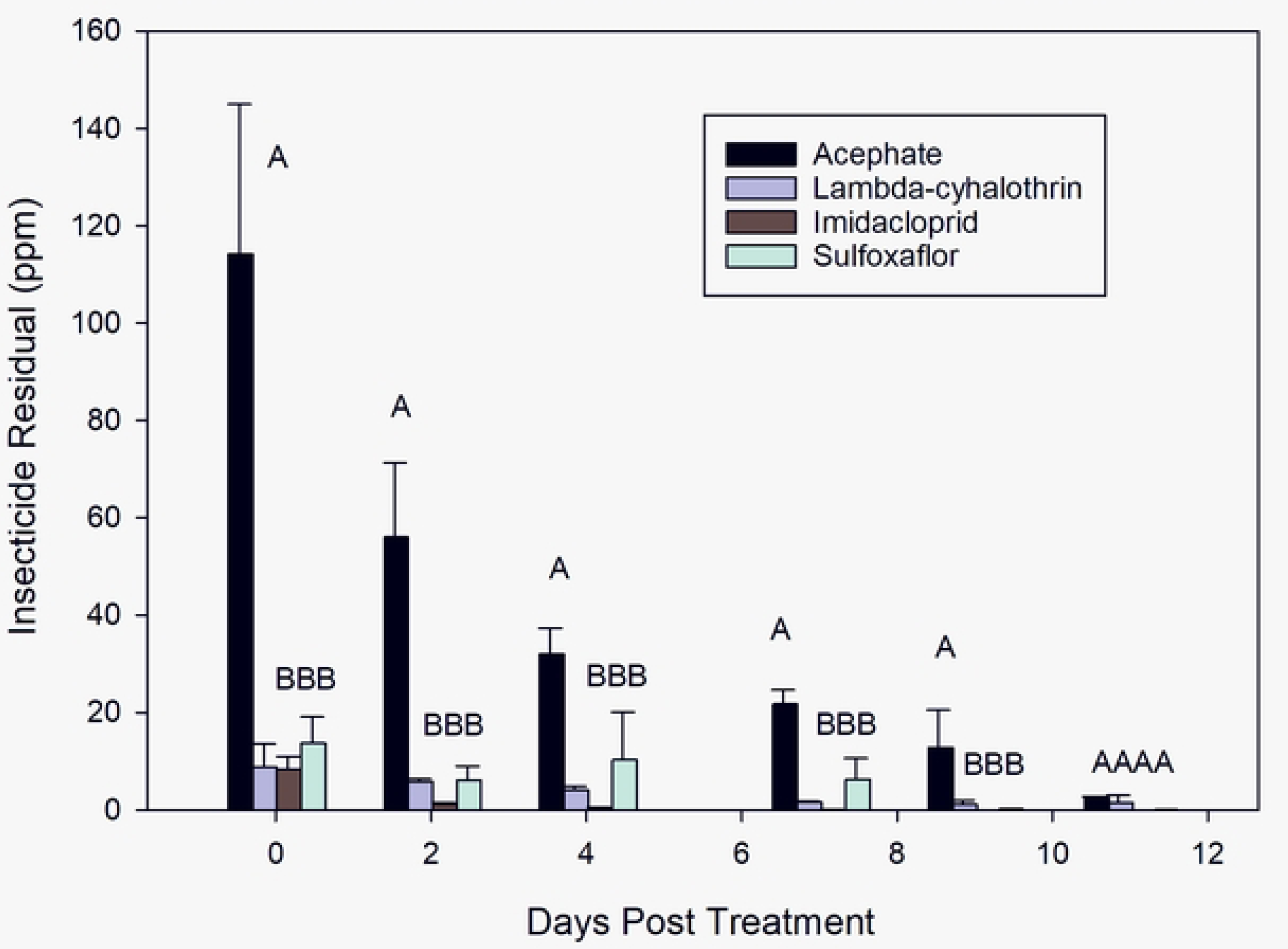
Insecticidal residual test of four classes of insecticides on leaf samples pulled at 0, 2, 4-, 7-, 9-, and 11-days post treatment periods. (Tukey’ HSD test, *p* = 0.05).

## Discussion

All insecticides tested were greatly affected due to the high levels of precipitation obtained in 2017. Our results demonstrated that environmental conditions may greatly influence persistence and degradation of foliar insecticides residues. Rainfall, especially when it occurs shortly after application can lead to immediate reduction of insecticide residues [41, 42]. Therefore. No leaf samples were collected after 7-DPT in 2017 for any treatment.

Thorough knowledge of the impacts related to contact toxicity of insecticidal residues is necessary for successful integrated pest management (IPM). Contact toxicity of residues is composed of two components including initial mortality experienced soon after application and the rate of changes in mortality as residues dissipate [43]. Our results demonstrated that the high concentration of all insecticides had the ability to remain present (ppm) till 11-DPT (Fig 5); however, no insecticide produced 100% cumulative mortality by contact even using a susceptible laboratory colony that has been successively reared for over 20 years with no infusion of field population (Figs 1 – 4). The organophosphate (acephate) showed greater toxicity than any other insecticide tested on *L. lineolaris* adults, with initial mortality of >50 % for 2016 and > 30 % for 2017 at 0-DAE on leaves pulled at 0-DPT for the high concentration. Mortality rapidly increased to 81.7 ± 23.4 and 63.3 ± 28.8 (SE) 1-day after exposure (DAE) for 2016 and 2017, respectively attaining 94.5 ± 9.5 and 95.4 ± 7.6 6-DAE for each year. This high residual activity persisted to 4-DPT with mortality >90% decreasing to >80% after 7-DPT in 2016. However, it is important to clarify, that the high mortality observed during this study is not likely to occur in field populations of *L. lineolaris* due to the current levels of resistance to acephate previously reported for this insect throughout the Mississippi Delta cropping region [1, 6–13]. For example, an earlier study demonstrated that acephate plus bifenthrine (a pyrethroid), bifenthrine alone, sulfoxaflor, dicrotophos (an organophosphate), and thiamethoxam (a neonicotinoid) all provided better control than acephate alone [44]. Yet, acephate and dicrotophos are two of the most used insecticides to control large populations of *L. lineolaris* in cotton with acephate being used more frequently [45].

Persistence is one of the main characteristics of residual activity, described in half-life, which is a comparative measure of the time needed for the chemical to degrade [46]. Therefore, the longer the insecticide’s half-life, the slower the degradation, which likely translate to increased persistence of the insecticide. In some cases, persistent insecticide residues are desirable due to its long-term control and the reduction in the need for multiple applications of the insecticide in high pressure situations. Our results demonstrated that the pyrethroid lambda-cyhalothrin could be as persistent insecticide as acephate. Interestingly acephate’s insecticidal residues (114.27 ppm) were >13.3 – fold higher than the pyrethroid (lambda cyhalothrin) (8.84 ppm) (Fig 5), yet the cumulative mortality attained with pyrethroid was almost as high as it was with acephate for both years (Figs 1 and 2). However, the initial mortality with acephate was 5-fold higher than lambda-cyhalothrin on the leaf sample pulled at 0-DPT in 2016. Similar initial mortality was observed between both insecticides in 2017, and cumulative mortalities were comparable for both insecticides at all evaluation periods; although, it increased faster with acephate than with lambda-cyhalothrin. This could explain why acephate is the most desirable control option to cotton growers, not just for the visible and faster control but for the reduction in needed insecticide applications [45]. Conversely, our results corroborated with Palmquist’ et al. 2012 [46] findings, which reported that pyrethroid insecticides as a group replaced many organophosphate insecticides due to better selectivity on target pest and less persistence than organochlorine insecticides. William et al. 2003 [43] demonstrated that the survival of female *Anaphes iole* (Hymenoptera: Mymiridae) was four times longer when exposed to the pyrethroid lambda-cyhalothrin than to the organophosphate acephate or the nicotinoid imidacloprid.

Imidacloprid, which is the world’s leading insecticide, was released in 2003 and has been approved for controlling infectious disease vectors [47]. Since then, it has been successfully used as a systemic and contact insecticide to control *L. lineolaris*, alone or in combination with other insecticides [3]. *L. lineolaris’* resistance to this insecticide it was first reported for the first time in 2012 [12], which was subsequently followed by several additional papers demonstrating the low response of this insect to neonicotinoids [13, 15, 34, 48, 49]. Our results confirmed their findings; however, our outcomes cannot be referred as a resistance, since our observations were limited to a susceptible laboratory colony established in 1998 with not infusion from field population [15]. The response of *L. lineolaris* to imidacloprid and its residual activity found in our study was significantly lower compared to acephate and lambda-cyhalothrin during 2016 and 2017 (Figs 1, 2 and 3). Although its insecticidal residues for the high concentration (8.35 ppm) on leaf samples collected at 0-DPT was as high as the pyrethroid insecticide (8.84 ppm) (Fig 5), its cumulative mortality did not exceed 40% and its systematic effect was delayed; indicating the effectiveness of contact insecticides over those systemics in nature [11]. The low cumulative mortality attained after 2-DPT period is supported by the low insecticidal residues found on the leaf samples tested. The ppm values decreased 6.62-fold from 0-DPT (8.35 ppm) to 2-DPT (1.27 ppm) and 596-fold to 9-DPT (0.014 ppm). These results are comparable to a recent study, that evaluated eight insecticides including imidacloprid, finding that this neonicotinoid had the lowest efficacy against *L. lineolaris* in cotton [49]. They reported that all insecticide treatments significantly reduced the *L. lineolaris* nymph populations; mortality due to imidacloprid, however, did not differ among untreated controls 14 days after treatment. Additionally, they noted that all insecticides treatments resulted in significantly higher yields than the untreated control, but imidacloprid treated plots had the lowest yield among treatments. Likewise, Graham and Smith, 2021 [50] corroborated with our finding, where they reported no significant differences among all treatments (insecticides and untreated control) seven days after treatment, where the total populations (nymph and adult) treated with imidacloprid were larger than untreated control.

Similar to imidacloprid, sulfoxaflor is a systemic insecticide that incurs its toxicity through contact and oral ingestion [50]. Therefore, based on *L. lineolaris* food consumption their systemic effect could be equal on nymphs and adults; however, the contact action could be higher in nymphs because they are less mobile than adults [15]. In this study, the mobility of adults was restrained, where their arena was a folded-treated cotton leaf in a 30 mL cup. Therefore, adults were forced to be in contact and feed on the cotton treated leaf. Our results demonstrated that efficacy of sulfoxaflor on *L. lineolaris* adults was highly superior to imidacloprid. The cumulative mortality trend for leaves pulled at 0-DPT was comparable to the pyrethroid lambda-cyhalothrin (Figs 3 and 4). Yet, the residual effect was as low as imidacloprid for leaf samples collected at 4, 7, and 9-DPT. The low cumulative mortality after the 4-DPT was unexpected, since the insecticidal residues test for sulfoxaflor was 2.18- and 3.82-fold higher than lambda-cyhalothrin for leaves pulled at 4-, and 7-DPT, respectively. These results deferred from other field studies using eight insecticides including sulfoxaflor, which demonstrated that sulfoxaflor’s efficacy against *L. lineolaris* in cotton was higher than any other insecticide, however they did not differ when compared to novaluron (an effective insect growth regulator) or acephate [49]. The authors reported no significant differences among 4, 7, and 14 days after treatment, suggesting that sulfoxaflor’s insecticide residual continued to 14 days after application. No differences in residual effect were found between sulfoxaflor and acephate [49]. Comparably, Siebert et al. 2012 [52] demonstrated similar residual efficacy levels at evaluation intervals >6 days after application between sulfoxaflor and acephate against moderately tolerant populations of *L. lineolaris* in 12 mid-southern U.S. locations from 2008 through 2010. Although, both studies were conducted a decade a part, acephate continually proves to demonstrate highly efficacious for control of *L. lineolaris* relative to other classes of insecticides regardless of any inherent resistance development.

Today, sulfoxaflor is considered one of the most promising insecticides used to control *L. lineolaris*. Our results however, indicated that the organophosphate acephate and the pyrethroid lambda-cyhalothrin exhibited higher mortality and longer residual activity than the sulfoxaflor (a sulfoxamine) on treated leaves with the highest concentration. *L. lineolaris* adults are significantly more susceptible to contact than systemic insecticides and due to its residual effect, acephate could kill over 80% of the TPB population 7-DPT. Conversely, in agricultural situations, selection of insecticides for *L. lineolaris* control needs to be closely related to dynamic resistance levels while preserving natural enemies of insect pests and pollinators, both of which can directly increase crop yields.

## Acknowledgements

The work is supported by USDA-ARS Research Project# 6066-22000-090-00D-Insect Control and Resistance Management in Corn, Cotton, Sorghum, Soybean, and Sweet Potato, and Alternative Approaches to Tarnished Plant Bug Control in the Southern United States. The authors would like to thank Tabatha Nelson, Arnell Patterson, and Henry Winter ARS-USDA, Southern Insect Management Research Unit (SIMRU), Stoneville, for their assistance with insect rearing, laboratory assays and field leave collection. Owen Houston, Phil Powell, G. (USDA ARS Southern Insect Management Research Unit, Stoneville, MS) for their assistance with the field plots preparation. To Michael Huoni, student trainee RIMRU for his valuable help in data entry. Mention of trade names or commercial products in this publication is solely for the purpose of providing specific information and does not imply recommendation or endorsement by the U.S. Department of Agriculture or the Agricultural Research Service.

## References

1. Maniania NK, Portilla M, Amnulla FM, Mfuti DK, Darie A, Dhiman G, Rao IM. Infectivity of Entomopathogenic Fungal Isolates Against Tarnished Plant Bug *Lygus lineolaris* (Hemiptera: Miridae). Journal of Insect Science. 2022;22(4):1–7. doi: 10.1093/jisesa/ieac040.

2. Gorge J, Glover JP, Gore J, Crow WD, Reddy GVP. Biology, Ecology, and Pest management of the Tarnished Plant Bug, *Lygus lineolaris* (Palisot de Beauvois) in Southern Row Crops. Insects. 2021;12(807):1–30. doi: 10.3390/insects12090807.

3. Mann RT. Seasonal management strategies for tarnished plant bug, *Lygus lineolaris* (Palisot de Beauvois), in midsouth cotton production systems. M.S. thesis. Mississippi State University, Mississippi State, MS. 2021. 56 p.

4. Snodgrass GL, Scott WP. Seasonal changes in pyrethroid resistance in tarnished plant bug (Heteroptera: Miridae) populations during a three-years period in the Delta of Arkansas, Louisiana, and Mississippi. Journal of Economic Entomology. 2000;93(2):441–446. doi: 10.1603/0022-0493-93.2.441.

5. Snodgrass GL, Gore J, Abel CA, Jackson R. Predicting field control of tarnished plant bug (Hemiptera: Miridae) populations with pyrethroid insecticides by use of glass-vial bioassays. Southwestern Entomologist. 2008;33(3):181–189.

6. Hollingsworth RG, Steinkraus DC, Tugwell NP. Responses of Arkansas populations of tarnished plant bugs (Heteroptera: Miridae) to insecticides, and tolerance differences between nymphs and adults. Journal of Economic Entomology. 1997;90(1):21–26. doi: 10.1093/jee/90.1.21.

7. Snodgrass GL. Pyrethroid resistance in a field population of the tarnished plant bug (Heteroptera: Miridae) in cotton in the Mississippi Delta. pp. 1186–1187. In Proceedings of the 1994 Beltwide Cotton Conference. 1994. National Cotton Council, Memphis, TN.

8. Snodgrass GL, Elzen GW. Insecticide resistance in a tarnished plant bug population in cotton in the Mississippi Delta. Southwestern Entomologist. 1995;20(3):317–232.

9. Snodgrass GL. Insecticide resistance in field populations of the tarnished plant bug (Heteroptera: Miridae) in cotton in the Mississippi Delta. Journal of Economic Entomology.1996;89(4):783–790. doi: 10.1093/jee/89.4.783.

10. Snodgrass GL. Glass-vial bioassay to estimate insecticide resistance in adult tarnished plan bugs (Heteroptera: Miridae). Journal of Economic Entomology. 1996;89(5):1053–1059. doi: 10.1093/jee/89.5.1053.

11. Snodgrass GL, Gore J, Abel CA, Jackson R. Acephate resistance in populations of the tarnished plant bug (Heteroptera: Miridae) from the Mississippi River Delta. Journal of Economic Entomology. 2009;102(2):699–707. doi: 10.1603/029.102.0231.

12. Allen KC, Jackson RE, Snodgrass GL, Musser FR. Comparative susceptibility of different life stages of the tarnished plant bug (Hemiptera: Miridae) to three classes of insecticide. Southwestern Entomology. 2012;37(3):271–280. doi: 10.3958/059.037.0303.

13. Parys KA, Luttrell RG, Snodgrass GL, Portilla M. Patterns of Tarnished Plant Bug (Hemiptera: Miridae) Resistance to Pyrethroid Insecticides in the lower Mississippi Delta for 2008-2015: Linkage to Pyrethroid use and cotton insect management. 2018;18(2):1–19. doi: 10.1093/jisesa/iey015.

14. Portilla M, Luttrell RG, Parys KA, Little N, Allen KC. Comparison of three bioassays methods to estimate levels of tarnished plant bug (Hemiptera: Miridae) susceptibility to acephate, imidacloprid, permethrin, sulfoxaflor, and thiomethoxam. Journal of Economic Entomology. 2018;111(6):2299–2808.

15. Portilla M. A laboratory Diet-overlay bioassay to monitor resistance in *Lygus lineolaris* (Hemiptera: Miridae) to insecticides commonly used in the Mississippi Delta. Journal of Insect Science. 2020;20(4):1–13. doi: 10.1093/jisesa/ieaa067.

16. Georghiou GP, Mellon RB. Pesticide Resistance in Time and Space. *In*: Georghiou GP and Saito T. (Ed.) Pest Resistance to Pesticide. pp. 1–47. Plenum Press. 1983. New York, NY. 797 pp.

17. North JH, Gore J, Catchot AL, Cook DR, Dodds DM, Musser FR. Quantifying the impact of excluding insecticide classes from cotton integrated pest management program in the U.U. Mid-South. Journal of Economic Entomology. 2019;112(1):341–348. doi: 10.1093/jee/toy339.

18. Cook DR, Threet M, Huff K. Cotton Insects Losses 2022. Mississippi State University Extension. Available online: https://www.biochemistry.msstate.edu/resources/2022loss.php (accessed on 07/10/2023).

19. Layton MB. Biology and damage of the tarnished plant bug, *Lygus lineolaris*, in cotton. Southwestern Entomologist. 2000;(23):7–20.

20. Musser FR, Stewart S, Bagwell R, Lorenz G, Catchot A, Burris E, Cool D, Robbins J, Greene J, Studebaker, Gore J. Comparison of direct and indirect sampling methods for tarnished plant bug (Hemiptera: Miridae) in flowering cotton. Journal of Economic Entomology. 2007;100(6):1916–1923. doi: 10.1093/jee/100.6.1916.

21. Wood W, Gore J, Catchot A, Cook D, Doods D, Krutz LJ. Susceptibility of Flowering cotton to damage and yield loss from tarnished plant bug (Hemiptera: Meridae). Journal of Economic Entomology. 2016;109(3):1188–1195. doi: 10.1093/jee/tow076.

22. Williams MR. Cotton confproc losses - 2016, pp. 710–754. In Proceeding Beltwide Cotton Conference, 4-6 Junauary 2017, Dallas, TX. National Cotton Council America, Memphis, TN.

23. Cook DR, Threet M. Cotton insect losses - 2021, pp. 145–200. In Proceeding Beltwide Cotton Conference, 4-6 January 2022, San Antonio, TX. National Cotton Council of America, Memphis, TN.

24. FORBES. “Silenced Data” Means we do not know global impacts of cotton pesticides. 2021. Available from https://www.forbes.com/sites/brookerobertsislam/2021/12/06/silenced-data-means-we-dont-know-global-impacts-of-cotton-pesticides/?sh=19e79691668b (accessed on 07/20/2023)

25. Scott WP, Snodgrass GL. Response of tarnished plant bugs (Heteroptera: Miridae) to traps baited with virgin males or females. Southwestern Entomologist. 2000;25(2):101–108.

26. Snodgrass GL, Gore J, Abel CA, Jackson R. Predicting field control of tarnished plant bug (Hemiptera: Miridae) populations with pyrethroid insecticides by use of glass-vial bioassays. Southwestern Entomologist. 2008;33(3):181–189. doi: 10.3958/0147-1724-33.3.181.

27. Fleming DE, Krishnan N, Catchot AL, Musser FR. Susceptibility to insecticides and activities of glutathione *S*-transferase and esterase in populations of *Lygus lineolaris* (Hemiptera: Meridae) in Mississippi. Pest Management Science. 2015;72(8): 1595–1603. doi: 10.1002/ps.4193.

28. Zhu YC, Luttell R. Variation of acephate susceptibility and correlation with esterase and glutathione *S*-tranferase activities in field populations of the tarnished plant bug, *Lygus lineolaris*. Pesticide Biochemistry and Physiology. 2012;103(3):202–209. doi: 10.1016/j.pestbp.2012.05.005.

29. Zhu YC, Snodgrass GL, Chen MS. Comparative study on glutathione S-tranferase activity, cDNA, and gene expression between malathion susceptible and resistant strains of the tarnished plant bug, Lygus lineolaris. 2007;87(1):62–72. doi: 10.1016/j.pestbp.2006.06.002.

30. Zhu YC, Snodgrass GL, Chen MS. Enhanced esterase gene expression and activity in a malathion-resistant strain of the tarnished plant bug, *Lygus lineolaris*. Insect Biochemistry and Molecular Biology. 2004;34(11):1175–1186. doi: 10.1016/j.ibmb.2004.07.008.

31. Zhu YC, Luttrell R. Altered gene regulation and potential association with metabolic resistance development to imidacloprid in the tarnished plant bug, *Lygus lineolaris*. Pest Management Science. 2014;71(1):40–57. doi: 10.1002/ps.3761.

32. Zhu YC, Guo Z, He Y, Luttrell R. Microarray analysis of gene regulations and potential association with acephate-resistance and fitness cost in *Lygus lineolaris*. PlosONE. 2012; doi: 10.1371/journal.pone.0037586.

33. Zhu YC, Snodgrass GL. Cytochrome p450cyp6x1 cDNAs and mRNA expression levels in three strains of the tarnished plant bug Lygus lineolaris (Heteroptera: Miridae) having different susceptibilities to pyrethroid insecticide. Insect Molecular Biology. 2003;12(1):39–49. doi: 10.1046/j.1365-2583.2003.00385.x.

34. Dorman S, Gross AD, Musser FR, Catchot BD, Smith RH, Reisig DD, Reay-Jones FPF, Greene JK, Roberts PM, Taylor SV. Resistance monitoring to four insecticides and mechanisms of resistance in *Lygus lineolaris* Palisot de Beauvois (Hemiptera: Miridae) populations of southeastern USA cotton. Pest Management Science. 2020;76(12):3935–3944. doi: 10.1002/ps.5940.

35. Du Y, Zhu YC, Portilla M, Zhang M, Reddy GVP. The mechanisms of metabolic resistance to pyrethroids and neonicotinoids fade away without selection pressure in the tarnished plant bug Lygus lineolaris. Pest Management Science. 2023; doi: 10.1002/ps.7570.

36. Portilla M, Snodgrass GL, Streett D. Effect of modification of the NI artificial diet on the biological fitness parameters of mass reared wester tarnished plant bug, *Lygus hesperus*. Journal of Insect Science. 2011;11(1):1–10. doi: 10.1673/031.011.14901

37. Cohen AC. New oligidic production diet for *Lygus hesperus* (Knight)and *L. lineolaris* (Palisot de Beauvois). Journal of Entomological Science. 2000;35(3):301–310. doi: 10.18474/0749-8004-35.3.301.

38. Institute S. SAS Software. 9.4 ed. Cary, NC: SAS Institute; 2013.

39. Abbott WS. A method of computing the effectiveness of an insecticide. Journal of Economic Entomology. 1925;18(2):265–7.

40. Robertson JL, Jones MM, Olguin E, Alberts B. Bioassays with arthropods. Boca Raton, FL: CRC press; 2017. 193 p Elston RC. An analogue to Fieller’s theorem using Scheffé’s solution to the Fisher-Behrens problem. The American Statistician. 1969;23(1):26–8.

41. Gautam BK, Little BA, Taylor MD, Jacobs JL, Lovett WE, Holland RM, Sial AA. Effect of simulated rainfall on the effectiveness of insecticides against spotted wing drosophila in blueberries. Crop Protection. 2016;81:122–128. doi: 10.1016/j.cropo.2015.12.017.

42. Mashaya N. Effect of simulated rain on efficacy of insecticide deposit on tobaco. Crop Protection. 1993;12(1):55–58. doi: 10.1016/0261-2194(93)90020-J.

43. Williams L, Price LD, Manrique V. Toxicity of field-weathered insecticide residues to *Anaphes iole* (Hymenoptera: Mymaridae), an egg parasitoid of *Lygus leneolaris* (Heteroptera: Miridae), and implications for inundative biological control in cotton. Biological Control. 2003;26(3):217–223. doi: 10.1016/S1049-9644(02)00157-3.

44. Graham SH, Catchot AL, Gore J, Cook DR, Doods D. Tarnished plant bug (Heteroptera: Miridae) behavioral responses to chemical insecticides. Insects. 2021;12(2):1072. doi: 10.3390/insects12121072.

45. Smith J, Crow WD, Catchot AL, Gore J, Cook DR, Musser F, Stewart SD, Brown S, Thrash B, Lorenz G, Batemen N, Studebaker G, Towles T, Kerns D. Evaluating efficacy and chemical concentrations of common unused insecticides targeting tarnished plant bug in Mid-South cotton. The Journal of Cotton Science. 2023;27:74–80.

46. Fishel FM. Pesticide Characteristics. IFAS Extension University of Florida. 2008;PI-166:1–3. Available online: https://edis.ifas.ufl.edu/publication/PI202 (accessed 06/20/2023).

47. Palmquist K, Salatas J, Fairbrother A. Pyrethroid insecticide: Use, environmental fate, and ecotoxicology. In: Perveen FK (ed.) Insecticide Advances in Integrated Pest Management. 2012; pp. 258–278. Intech Press, Croatia. 708 pp.

48. Parys KA, Luttrell RG, Snodgrass GL, Portilla M, Copes JT. Longitudinal measurement of tarnished plant bug (Hemiptera: Miridae) susceptibility to insecticides in Arkansas, Louisiana, and Mississippi: Associations with insecticide use and insect control recommendations. Insects. 2017;109(8):4–21. doi: 10.3390/insects8040109.

49. Mississippi State University Extension. Insecticide Performance on Tarnished Plant Bugs in Mississippi Cotton. Publication 3548 (POD-10-20) 2023. Available online: http://extension.msstate.edu/publications/insecticide-performance-tarnished-plant-bugs-mississippi-cotton (Accessed 07/13/2023).

50. Graham SH, Smith RH. Evaluation of insecticide for control of tarnished plant bugs in North Alabama cotton, 2020. Arthropod Management Tests. 2021;46(1):1–1. doi: 10.1093/amt/tsab090.

51. Zhu YC, Yao J, Adamczyk, Luttrell R. Synergistic toxicity and physiological impact of imidacloprid alone and binary mixtures with seven representative pesticides on honey bee (Apis mellifera). PlosONE. 2017;12(5):e0176837. doi: 10.1371/journal.pone.0176837.

52. Siebert MW, Thomas JD, Nolting SP, Leonard BR, Gore J, Catchot A, Lorenz GM, Stewart DS, Cook DR, Walton LC, Lassiter RB, Haygood RA. Field evaluation of sulfoxaflor, a novel insecticide, against tarnished plant bug (Hemiptera: Miridae) in cotton. Journal of Cotton Science. 2012;12(2):129–143.

